# Potassium isomerized linoleate satisfies EPA MB-35-00 for surface disinfection of *Candida auris* (AR-0381)

**DOI:** 10.1101/2021.02.23.432518

**Authors:** David Changaris, Anne Carenbauer

## Abstract

Four distinct lots of Potassium Isomerized Linoleate, 442 mM or 141 mg/ml (within the cosmetic-cleanser and commercial-soap acceptable concentration ranges but much greater than physiologic conditions), showed greater than 5 log kill rates (total) for *Candida auris* (AR-0381) for all carriers during 5 separate procedural runs of EPA MB-35-00. After exposing the inoculated stainless-steel carriers to the plant-oil soap for 1, 2, or 10 minute(s), we recovered no *Candida auris* on these tested carriers. Control carriers with phosphate-buffered saline applied, recovered 5.0-6.8 Log of colony-forming units per carrier.

## Introduction

The United States Center for Disease Control and Prevention (CDC) identifies *C. auris* as a major global threat, with fatality rates in immunocompromised individuals ranging from 10-60% **[1]**. Finding safe and effective non-toxic disinfectants against *C. auris* has proven difficult **[2]**, **[3]**, **[4]**, **[5]**, **[6]**. The plant oil Isomerized Linoleic acid has maintained Generally Recognized as Safe (GRAS) status as a food additive since 2008 **[7]**. Making a potassium salt of a plant oil remains a simple and readily achievable process to effect a cosmetically acceptable cleanser or soap. Current regulatory guidance from the Environmental Protection Agency (EPA) states: “In 1988, the EPA determined that soap salts have ‘no independent pesticidal activity’ in antimicrobial products, and must be classified as inert ingredients in those products … Antimicrobials that still contain soap salts as active ingredients are considered misbranded….“”**[8]**

Linoleic acid has numerous reported anti-fungal effects, including isomerized linoleic acid known to reduce hyphae formation in *C. albicans* **[9]**(Shareck et al., 2011). Methyl esters of sunflower oils appear to inhibit *Candida glabrata*. And laurel seed oil and coriander seed oil show antifungal effects against the fungi type *C. albicans* **[10]**(Pinto et al., 2017). Lauric acid, myristoleic acid, linoleic acid and arachidonic acid derived from bovine whey inhibits the germination of *C. albicans* **[11]**(Cansel, 2019). Recent reports link quorum sensing Farnesol like activity to 31 saturated and unsaturated fatty acids, six medium-chain saturated fatty acids, that is, heptanoic acid, octanoic acid, nonanoic acid, decanoic acid, undecanoic acid, and lauric acid, effectively inhibited *C. albicans* biofilm formation by more than 75% at 2 μg ml-1 with MICs in the range 100–200 μg ml-1**[12]**(Lee et al., 2020). The many natural sources of anti-*C. albicans* compounds include “traditional Chinese medicine”**[13]**(Gong et al., 2019). The concentration used herein, 442 mM or 141 mg/ml, falls within the range of plant oil salts generally used as commercial cosmetic cleansers or soaps **[14][15]**(Thompson, 2014, Ade, 2020).

## Objective

To test the biocidal capacity of potassium isomerized linoleate against *C. auris* (AR-0381) using a standard biocidal test, US Environmental Protection Agency protocol MB-35-00.

## Results & Discussion

Four distinct lots of Potassium Isomerized Linoleate, 442 mM, showed greater than 5 log kill rates (total) for *C. auris* (AR-0381) for all carriers during 5 separate procedural runs of SOP EPA MB-35-00 **[16]**. All aspects of the SOP were applied, including: Pegen laser-cut carriers; generation of −80°C glycerol stocks; pre-application of the “soil load” containing mucin, bovine serum albumin, and yeast extract; and drying the carriers under vacuum. We processed these carriers for testing the same-day we made them. Exposing the 10 mm *C. auris*-inoculated and dried stainless steel carriers to the soap salt at 1, 2, and 10 min showed no *C. auris* recovery on the membrane filters after 120 hours of incubation on Sabourad-dextrose (Emmons) agar at 31°C. Phosphate-buffered saline controls of the same carriers provided consistent results between 5.0-6.8 log colony forming units per carrier, when inputs of culture were controlled using OD_600_ between 15-19, in phosphate-buffered saline. Dilutions of 1 to 10 of the culture were used to maintain range within the UltraSpec Cell Monitor.

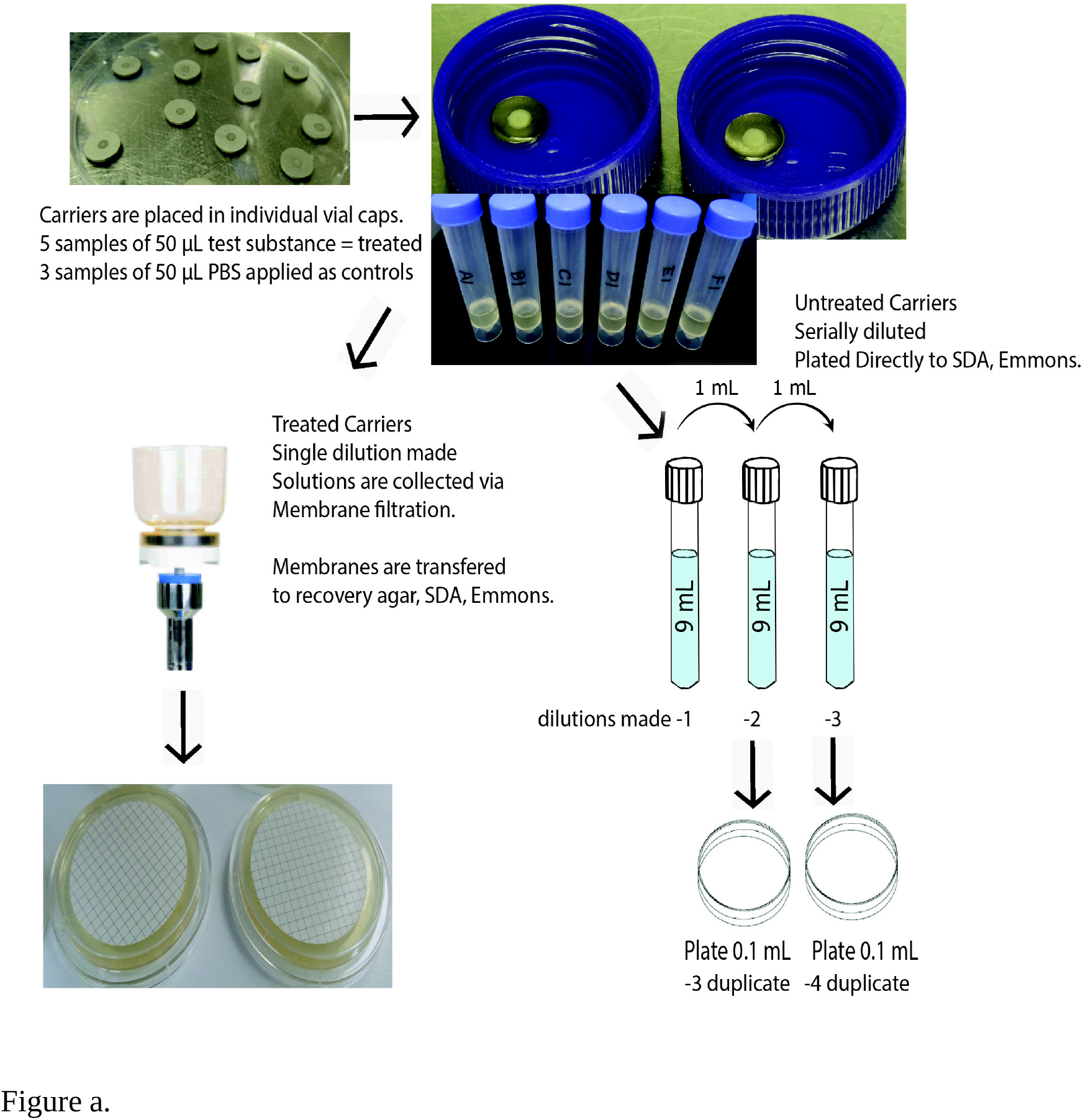
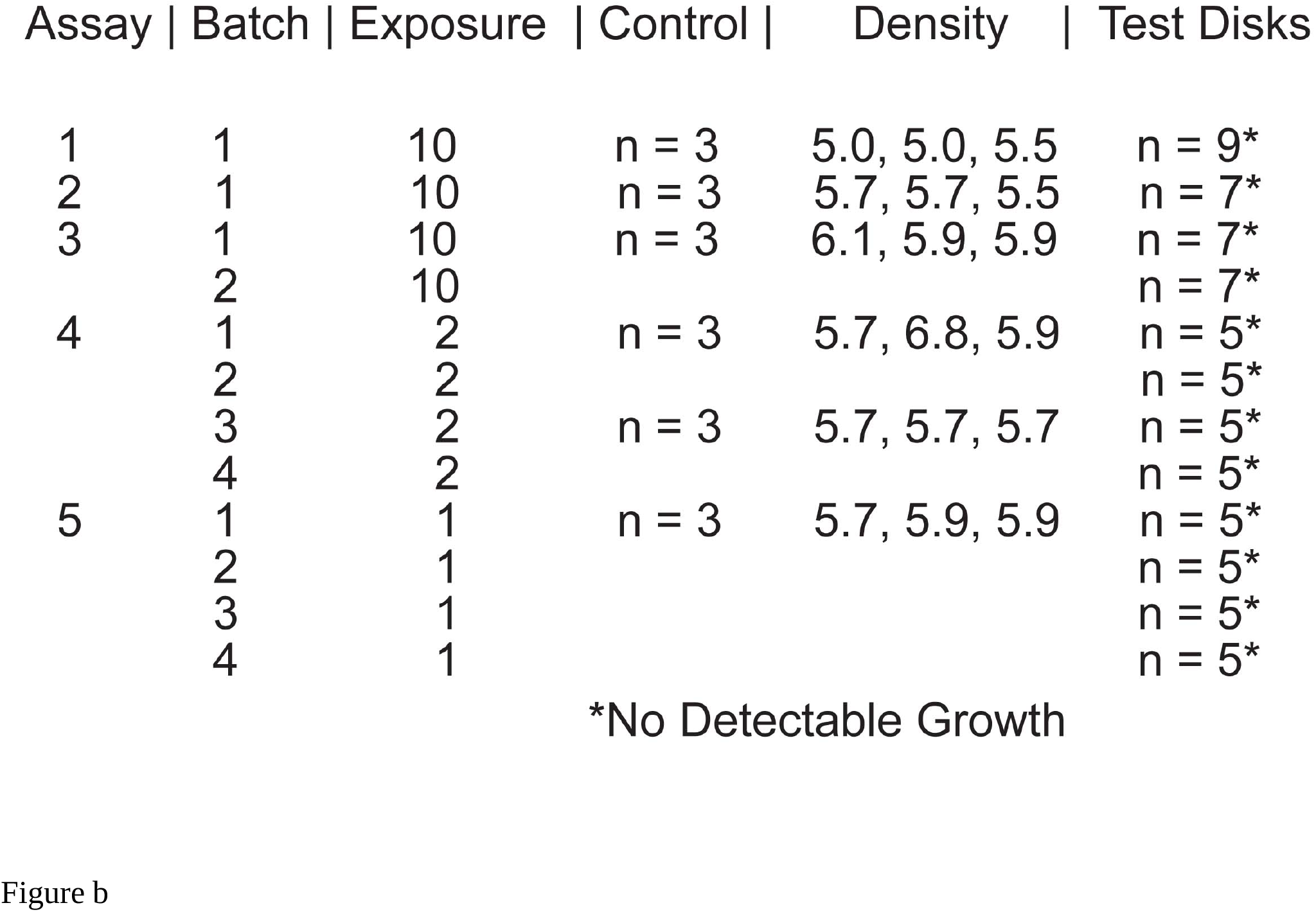
US EPA Protocol MB-35-00 (a) run on five separate dates testing the efficacy of potassium isomerized linoleate (442 mM or 141 mg/ml) against Candida auris (AR-0381) documented greater than 5 log suppression with exposures at 1, 2, and 10 minutes (b). “SDA” references Saboraud dextrose agar. The EPA protocol establishes the procedure by which one satisfies the requirement of a 5-log reduction for certification. Before inoculation, we estimated by O.D. 600, that each assay parent culture of C. auris contained between 5-6 log density. The column labeled “Density” represents the “control” C. auris densities by plating (b). For example, Assay 1 reports “5.0, 5.0, 5.5.” This means that within the “Control” part of Assay 1, we inoculated three separate “carries” with C. auris. We washed each “carry” with the neutralization buffer, putting each through serial dilution in PBS and plating in duplicate for counts of CFU/mL. We obtained the values of 5.0 log for two “carries” and 5.5 logs for the remaining “carry.” We processed 18 “Control Carries” with cultured-plate densities of 5-6 logs, in duplicate distributed over 5 assays. For the “Experimental” or “Treated Arm” at 10-minute exposure, 30 “carries” were inoculated and subsequently treated and cultured, showing no growth of C.

## Conclusions

In summary, we have shown that a plant oil soap salt, potassium isomerized linoleate has the germicidal capacity to effectively disinfect *C. auris* (AR-0381) on hard non-porous surfaces within US EPA Protocol MB-35-00.

## Limitations

The mechanism of potassium isomerized linoleate biocidal activity against *C. auris* (AR-0381) remains obscure.

## Alternative Explanations

The plant-oil soap may lyse the cells’ walls, bind to cell membranes and ion channel proteins, or alter essential functions.

## Conjectures

The central conjugated-pi bonds within the lipid may attract molecular complexes on the surface structures **[17]**, **[18]**. The alpha-carboxyl salt may interact as a detergent with local hydrophilic structures while the omega aliphatic end may disorder hydrophobic surface structures. Distinct amino acid residues of the membrane ion channels may also provide Diels-Alder-adduct-affinity sites inside the channels, facilitating disruption through osmotic lysis. Signaling lipids of non-conjugated families are known to diffuse through membranes to affect nuclear mechanisms in yeast and bacteria **[19]**, **[20]**, **[21]**.

We speculate that the 442 mM or 141 mg/ml concentration is close to minimal concentration to successfully fulfill the 1 min exposure using US EPA Protocol MB-35-00.

## Methods

We followed the United States Environmental Protection Agency Protocol MB-35-00 as written **[16]**. We dispersed *Candida auris* (AR-0381) obtained from the CDC AR Isolate Bank into glycerol stocks and stored at −80°C.

Each experiment began with an 8 mL overnight culture in Sabouraud dextrose broth (Emmons) from fresh −80°C glycerol stock. The defined procedure included harvesting by centrifugation, re-suspension in phosphate-buffered saline, and mixing with a 3-part soil load containing both with yeast extract powder (RM027, Himedia), bovine serum albumin (A2153, Sigma Life Science), and gastric mucin (HY-B2196/CS-7626, Med Chem Express). We inoculated the prepared 10 mm metal carriers (430-107L, Pegen Industries) and dried them in a desiccator under vacuum with a drying resin. For the treated group, we coated each disk with 50 μL 0.442 M Potassium Isomerized Linoleate onto the disk surface for 1, 2, or 10 minutes. Staggered starting times allowed for the full set of carriers. Carriers were transferred with sterile forceps to 10 mL neutralization solution (Sabouraud Dextrose Broth, Himedia), and vortexed for 1 minute. We filtered the entire neutralization solution plus 2 phosphate-buffered saline washes from the carriers, through membrane filters (200300-01, Rocker) with 47 mm PES 0.45 micron pore size (PES4547100, Sterlitech). We placed each membrane onto Sabouraud-dextrose-agar-(Emmons) plates and cultured for 120 hours at 31°C.

We processed the controls similarly, using phosphate-buffered saline (50 μL) applied to inoculated metal-disk carriers. We transferred these disks to 10 mL Sabouraud-dextrose-broth-neutralization solution, diluted serially in phosphate-buffered saline, and plated directly to Sabouraud-dextrose-agar (Emmons) plates to culture for 120 hours at 31°C. The MB-35-00 protocol identifies this method between treated and control samples to eliminate possible cross-contamination of the test samples in the filtration process.

## Funding Statement

After Accident Care, LLC provided the funds for this study.

## Acknowledgements

We thank the EPA and CDC for graciously providing the *Candida auris* strain AR-0381 for study.

## Conflict of interest

The authors do declare conflicts of interest: David G. Changaris is the sole owner of Ceela Naturals, LLC providing the potassium isomerized linoleate, After Accident Care, LLC providing funding for this study, and US patents concerning antimicrobial capacity of isomerized linoleic acid.

## Ethics Statement

No fraudulence was committed in performing these experiments or during processing of the data.

